# A frequency peak at 3.1 kHz obtained from the spectral analysis of the cochlear implant electrocochleography noise

**DOI:** 10.1101/2023.09.09.556985

**Authors:** Javiera Herrada, Vicente Medel, Constantino Dragicevic, Juan C. Maass, Carlos E. Stott, Paul H. Delano

## Abstract

**Introduction:** The functional evaluation of auditory-nerve activity in spontaneous conditions has remained elusive in humans. In animals, the frequency analysis of the round-window electrical noise recorded by means of electrocochleography yields a frequency peak at around 900 to 1000 Hz, which has been proposed to reflect auditory-nerve spontaneous activity. Here, we studied the spectral components of the electrical noise obtained from cochlear implant electrocochleography in humans.

**Methods:** We recruited adult cochlear implant recipients from the Clinical Hospital of the Universidad de Chile, between the years 2021 and 2022. We used the AIM System from Advanced Bionics® to obtain single trial electrocochleography signals from the most apical electrode in cochlear implant users. We performed a protocol to study spontaneous activity and auditory responses to 0.5 and 2 kHz tones

**Results:** Twenty subjects including 12 females, with a mean age of 57.9 ± 12.6 years (range between 36 and 78 years) were recruited. The electrical noise of the single trial cochlear implant electrocochleography signal yielded a reliable peak at 3.1 kHz in 55% of the cases (11 out of 20 subjects), while an oscillatory pattern that masked the spectrum was observed in seven cases. In the other two cases, the single-trial noise was not classifiable. Auditory stimulation at 0.5 kHz and 2.0 kHz did not change the amplitude of the 3.1 kHz frequency peak.

**Conclusion:** We found two main types of noise patterns in the frequency analysis of the single-trial noise from cochlear implant electrocochleography, including a peak at 3.1 kHz that might reflect auditory-nerve spontaneous activity, while the oscillatory pattern probably corresponds to an artifact.

## Introduction

The functionality of the auditory nerve is essential for transmitting afferent activity to the central auditory system, allowing sound perception and speech comprehension (Kiang 1965; Kiang et al., 1967; Louage et al., 2004). Several methods have been developed to study the auditory nerve function, such as the auditory-evoked compound action potentials obtained with electrocochleography (Galambos, 1956; Delano et al., 2007; Eggermont, 2019), electrically evoked compound action potentials (ECAPs) obtained with cochlear implants (CI) (Hey and Müller-Deile, 2015), wave I auditory brainstem responses (Steinhoff et al., 1988; Kujawa and Liberman, 2009), or auditory-nerve compound responses recorded during neurosurgery in humans (Yamakami et al., 2003; Ishikawa et al., 2017). However, these techniques need to evoke neural responses using auditory or electrical stimulation (Maggu, 2022), making it unfeasible to record auditory-nerve spontaneous activity.

In order to study auditory-nerve spontaneous activity, researchers have performed invasive recordings in animal models, including cats, guinea pigs and chinchillas (Kiang et al., 1976; Manley et al., 1976; Liberman, 1978; Temchin et al., 2008). These studies evaluated auditory-nerve single fiber activity, allowing the waveform characterization of the spontaneous spiking activity, which is also known as the “unitary response” of auditory-nerve neurons. These waveforms last less than 1 ms in cats (Wang, 1979), and it is accepted that their additive responses to synchronizing stimuli, such as clicks or electrical pulses, constitute auditory-nerve composed responses (Wang, 1979; Dolan et al., 1983), such as ECAPs (Hey and Müller-Deile, 2015). However, to our knowledge, the spontaneous activity of auditory-nerve single fibers in humans has never been reported (Dong et al., 2020; Dong et al., 2021; Dong et al., 2023).

An interesting approach, which has been used to study spontaneous auditory-nerve activity in animals, was introduced by Dolan et al. in 1990. These authors studied the electrical noise recorded with a round-window electrocochleography in guinea pigs, and reported a frequency peak around 900 Hz obtained from the spectrum of the spontaneous electrical activity. Later, Searchfield and colleagues (2004) showed that this peak at 900 Hz disappeared after blocking neural conduction with tetrodotoxin, suggesting a neural origin.

In humans, we used non-invasive tympanic membrane electrocochleography with wick electrodes, and found a reliable peak at 1 kHz in the frequency spectrum of the electrical noise recorded from the eardrum in silent conditions (Pardo-Jadue et al., 2017). However, the amplitude of this frequency peak was affected by stimuli of different modalities in a non-specific manner, as auditory and bithermal caloric vestibular stimulation could elicit responses, suggesting a mixed origin for this peak, not limited to the auditory modality, and probably including peripheral and central nervous structures as possible sources.

Cochlear implants are neuroprosthetic devices that have become the gold standard for the treatment of severe to profound hearing loss (Rauschecker and Shannon, 2002; Carlyon and Goehring 2021). In addition to the primary goal of restoring audition, cochlear implants can also be used for recording intracochlear electrocochleography, allowing the measurement of auditory-nerve ECAPs (Undurraga et al., 2010; Hey and Müller-Deile, 2015), or acoustically evoked cochlear microphonics (CM), as a measure of cochlear hair cell function (Giardina et al., 2019; O’Leary et al., 2023). However, a reliable method for recording the spontaneous activity of auditory nerve neurons in humans is still lacking (Dong et al., 2020; Dong et al., 2021; Dong et al., 2023).

Here, we propose to study the frequency components of the single-trial electrical noise recorded with cochlear implant electrocochleography in spontaneous and acoustically induced conditions in humans.

## Methods

### Subjects

Adult unilateral cochlear implant (HiRes, Advanced Bionics®) recipients from the Otolaryngology Department at the Hospital Clínico de la Universidad de Chile program, were recruited between the years 2021 and 2022. All procedures were approved by the Institutional ethical committee of the Hospital Clínico de la Universidad de Chile, and all volunteers signed an informed consent.

### Evaluations

All subjects were evaluated with a comprehensive battery of audiological tests, including unaided pure tone audiometry using headphones, free field audiometry, speech comprehension in silence and noise conditions in aided conditions with cochlear implants. Hearing thresholds, and free field hearing thresholds with cochlear implants were obtained using a clinical audiometer (AC40 Hybrid – Interacoustics®) at 0.25, 0.5, 1, 2, 3, 4, 6 and 8 kHz.

### Speech comprehension in quiet conditions

We used the International Matrix Test (Oldenburg Measurement Applications) to determine the minimum intensity (in dB) to obtain 50% of speech intelligibility presenting 30 sentences in quiet conditions.

### Speech in noise

We used the International Matrix Test (Oldenburg Measurement Applications) to determine the minimum signal to noise relation (SNR in dB) to obtain 50% of speech intelligibility, presenting 30 sentences in background broad-band noise conditions at 65 dB SPL.

### Electrocochleography

Postsurgery electrocochleography recordings were performed at least six months after cochlear implant surgery. We used the cochlear implant (HiRes, Slim J and MidScala) connected to an AIM System from Advanced Bionics® to obtain single-trial electrocochleography signals from the most apical electrode in cochlear implant recipients. We also performed an experimental control in ex-vivo conditions using a cochlear implant (HiRes, Slim J) immersed in 0.9% saline solution to record the input signal during the 9-minutes protocol (see below). Electrocochleography data was obtained at 9280 samples per second, while electrode impedances were ≤12 kΩ. We calculated the amplitude of cochlear microphonic responses to the 0.5 kHz and 2 kHz stimuli from the power spectral density (PSD) by using the median of the short-time Fourier Transform (STFT) using two different strategies: evoked and induced PSDs. Evoked spectrums were obtained by calculating the STFT over the averaged signal of all trials, while induced spectrums were obtained by the average of all single-trial STFT. In order to measure the baseline level of noise, we used spectral parametrization (Donoghue et al., 2020) to obtain broadband power and narrowband relative power measures using the fitting oscillations and one-over-F (FOOOF) algorithm.

### Auditory stimuli

We performed a nine-minute protocol to record the cochlear implant electrocochleography signal during spontaneous and auditory stimulation conditions (Figure 1). The protocol included three periods of three minutes each: (i) spontaneous activity in silent condition, (ii) evoked and induced responses at 0.5 and 2 kHz presented at 115 and 110 dB correspondingly, and (iii) post-stimulus spontaneous activity in silent conditions. Tones were presented using alternating phases (180° difference) in odd and even trials, with a duration of 50 ms per trial and on/off ramps of five ms. Each three minute period included a total of 3600 single-trials, including 1800 trials per phase in the period of acoustic stimulation. The complete 9-minute protocol included 10,800 trials. Electrocochleography recordings were performed in two different soundproof rooms of the Otolaryngology Department, located in the first and fourth floor of the Hospital. A digital microphone recorder TASCAM DR-05 (sampling rate: 44100 Hz, digitized at 16 bits) was used in control experiments to rule out a sound source at the 3.1 kHz frequency band.

**Figure 1.**
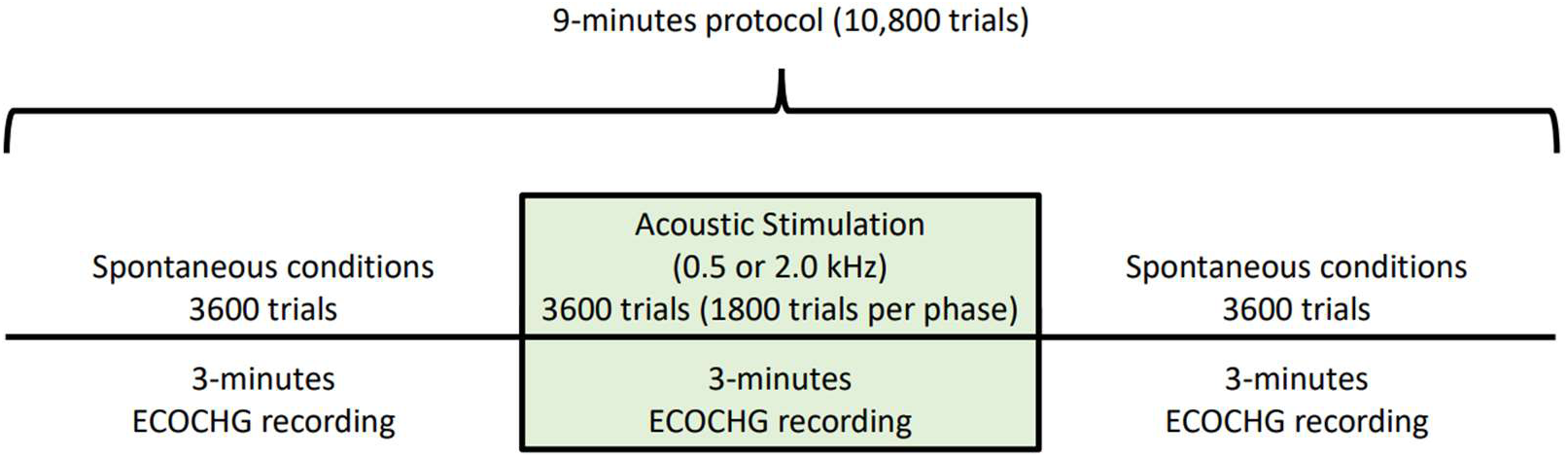
Timeline of the 9-minutes protocol. The protocol included three periods of cochlear implant electrocochleography (ECOCHG) recordings. The first and last three-minutes periods were acquired in spontaneous (silent) conditions, while the middle period included acoustic stimulation with 0.5 or 2 kHz tones. Each three-minutes period included 3600 trials. The 3600 trials of the stimulation period included 1800 trials of two 180° alternating polarity stimuli.

## Results

Measurements were obtained from twenty adult subjects (12 female, 57.9 ± 12.6 years (mean ± standard deviation), range between 36 and 78 years old; 14.4 ± 3.5 years of education (mean ± standard deviation)) with bilateral, severe to profound sensorineural hearing loss implanted with unilateral cochlear implants. A summary of demographic and audiological data is presented in Table 1. Regarding hearing thresholds, subjects had unaided pure tone averages (PTA) above 80 dB HL, while using cochlear implants, the average PTA was better than 50 dB HL (Figure 2).

**Figure 2.**
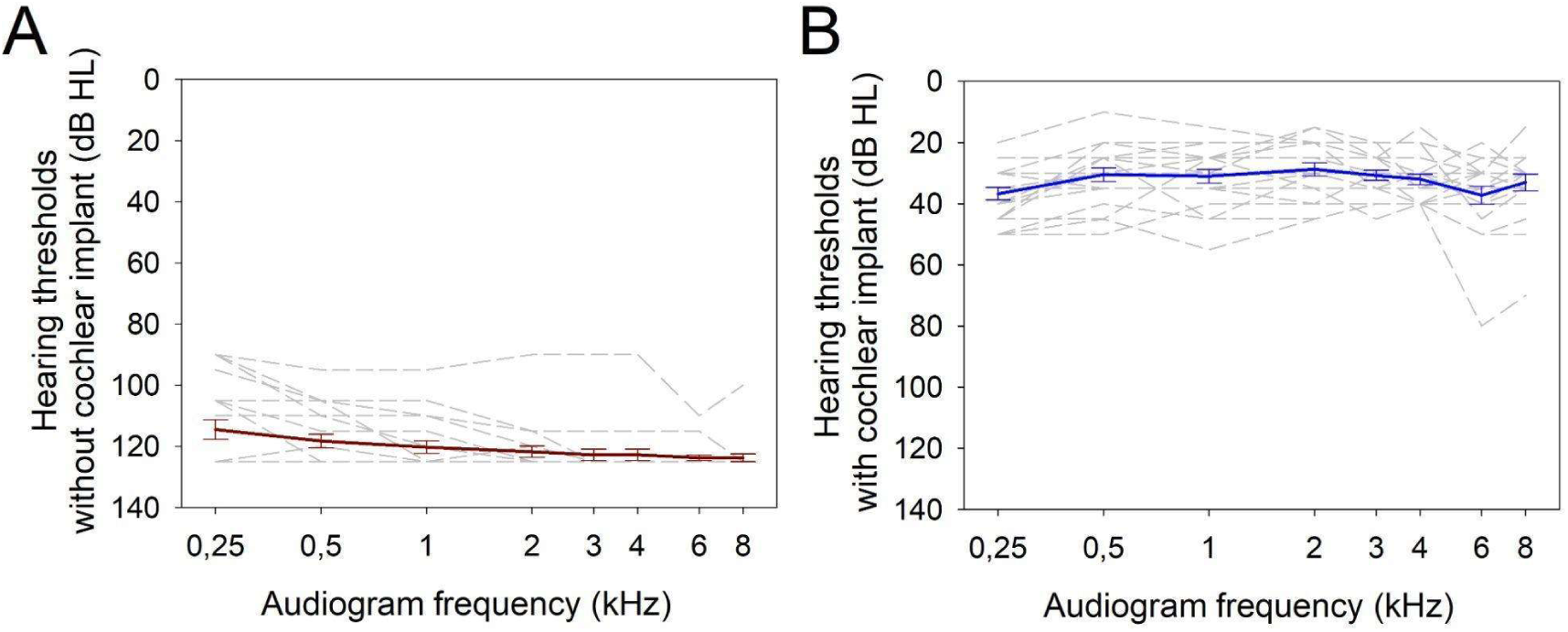
Audiogram thresholds (n=20). (A) Audiogram thresholds of the implanted ear without cochlear implants (red trace shows mean ± SEM). (B) Free field audiogram thresholds using cochlear implants (blue trace shows mean ± SEM). Segmented lines illustrate individual cases.

**Table 1.**
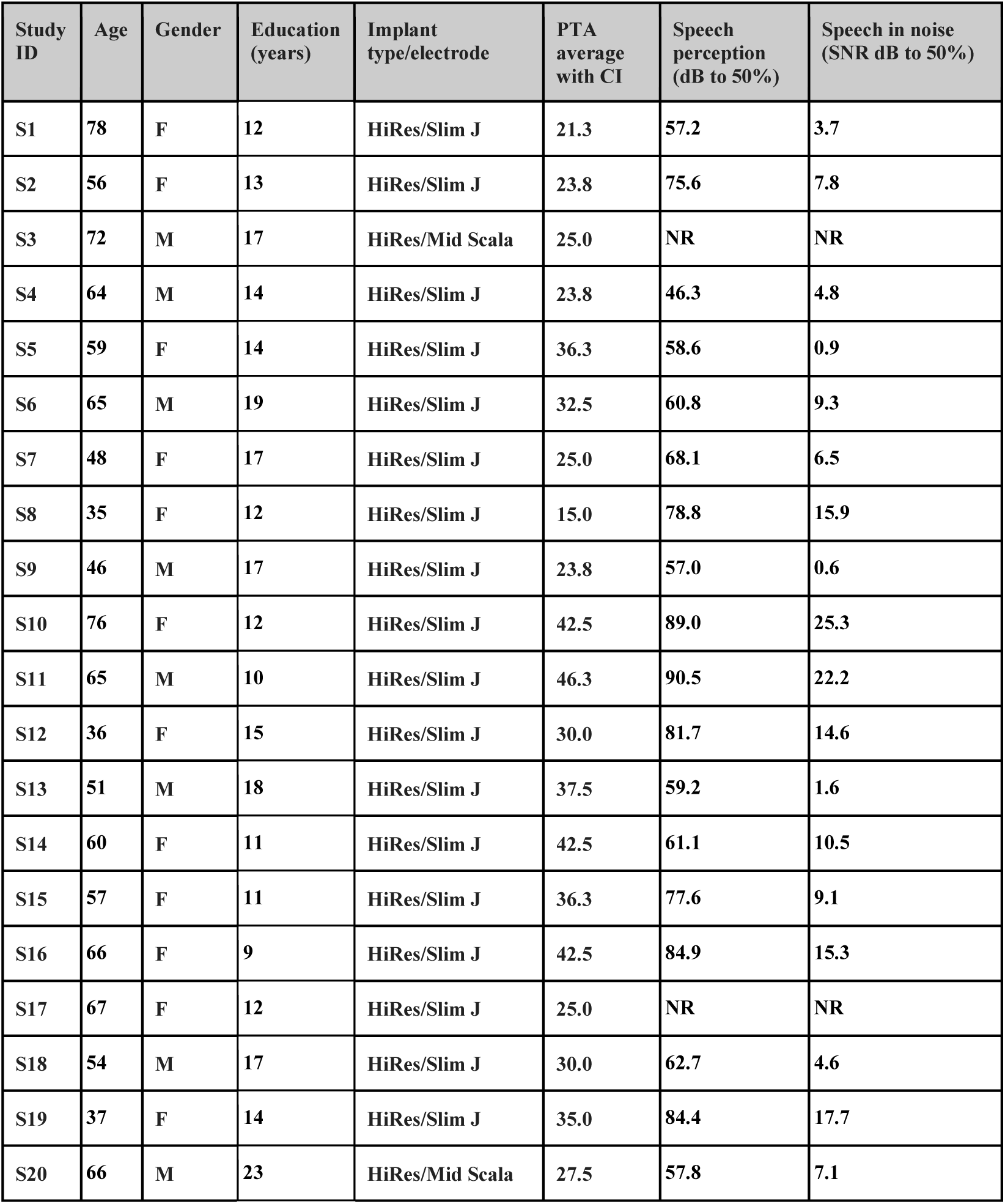
Summary of participants demographic and audiological data.

## Electrocochleography

We used two different methods to analyze the same electrocochleography data, including “evoked” and “induced” approaches (see methods section for more details), in spontaneous conditions (silence), and during auditory stimulation with a 0.5 and 2 kHz tone. Briefly, in the evoked condition, trials were averaged over time, and an STFT was computed for the averaged (evoked) signal, while in the induced analysis method, single-trial STFTs were calculated, and the STFT power spectrums of the 3600 trials (or 1800 trials of each phase) were averaged to obtain a grand average STFT of a given condition in the experimental protocol. These approaches gave us two STFT of the same data, evoked and induced, that we show and compare in the following results.

Figure 3 shows an example of the spectrograms obtained with the evoked and induced analyses of the cochlear implant electrocochleography signals in spontaneous (silence) conditions. Notice that in the induced condition a frequency peak at 3.1 kHz can be observed in the spectrum, and that the broadband power (background noise level) was greater when measured with induced techniques as compared to the evoked analysis (Figure 3). We found the frequency peak at 3.1 kHz in 55% of the cases (11 out of 20), while an oscillatory pattern that masked the 3.1 kHz frequency band in the spectrum was observed in seven cases (7 out of 20) (Figure 4). In addition, in two cases the single trial noise was not classifiable (not shown).

**Figure 3.**
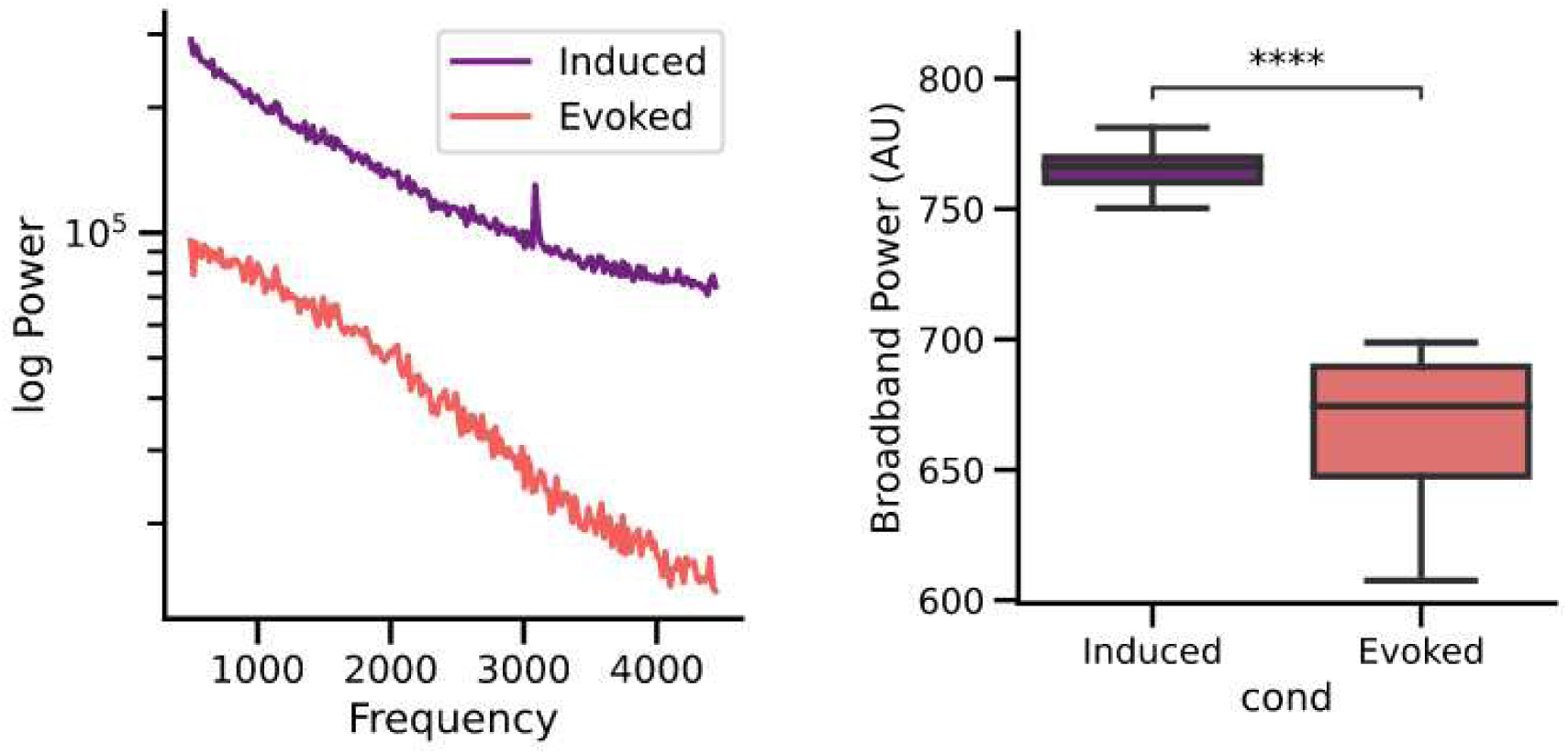
Spectrograms obtained with evoked (red trace) and induced (purple trace) analysis methods in spontaneous conditions (no auditory stimulation) for the same electrocochleography cochlear implant data. Notice the presence of a 3.1 kHz peak in the power spectrum of the induced, but not in the evoked analysis method. Box-plots (n=20) show that the broadband noise (total power under the curve) is larger in the induced condition as compared to the evoked method (****t-test paired samples p=2.34e-07) (AU: arbitrary units).

**Figure 4.**
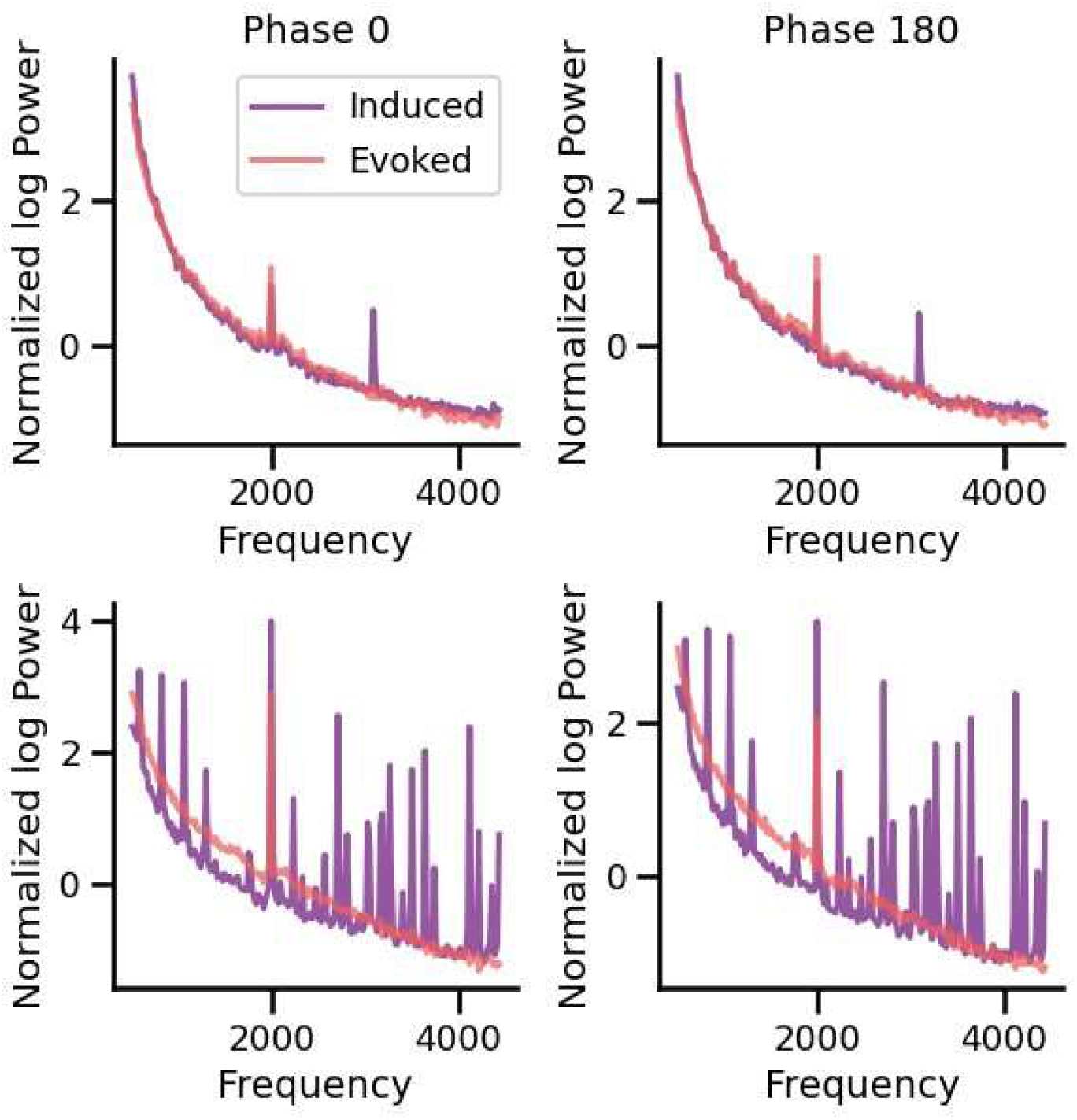
Power spectra of the cochlear implant noise obtained from two individuals (top and bottom panels) during the 2 kHz tone stimulation. Induced and evoked STFTs were normalized to the aperiodic 1/f baseline curve for better comparison of the amplitudes of the frequency peaks. Red spectra were calculated to the averaged waveform (evoked STFT), while purple spectra were computed averaging single trial STFT (induced STFT). The upper row illustrates an example of a case in which there was a peak at the 2 kHz cochlear microphonic response and at 3.1 kHz observed with both alternating stimuli (phase 0° and 180°) in induced conditions (purple STFT). If the same signal is averaged (evoked), and the STFT is computed to the averaged waveform, only the evoked 2 kHz cochlear microphonic response is present, while the 3.1 kHz peak is not found (red STFT). The bottom row shows an example obtained from another subject, in which the induced STFT yields an oscillatory pattern (purple spectrum) that masks the power spectrum, including the 3.1 kHz frequency band. On the other hand, in the evoked STFT a clear cochlear microphonic response at 2 kHz is visible.

Figure 4 shows examples of the 3.1 kHz frequency peak and the oscillatory pattern obtained during auditory stimulation with a 2 kHz tone. Cochlear microphonics responses were obtained with both types of analysis as a frequency peak at 0.5 or 2 kHz. We found that on average the amplitude of the 0.5 and 2.0 kHz cochlear microphonics responses were larger with evoked than with induced conditions (0.5 kHz: t-test, p=2.584e-4; 2 kHz: t-test, p=1.921e-4).

To test whether the 3.1 kHz frequency peak found with the single trial STFT induced method analysis is an artifact of the cochlear implant electronics, we recorded the same protocol using a cochlear implant immersed in saline solution outside a human being. Figure 5 shows that in the ex-vivo condition there was no frequency peak at 3.1 kHz, as compared to the spontaneous recordings made in silent conditions in a human patient with a cochlear implant.

**Figure 5.**
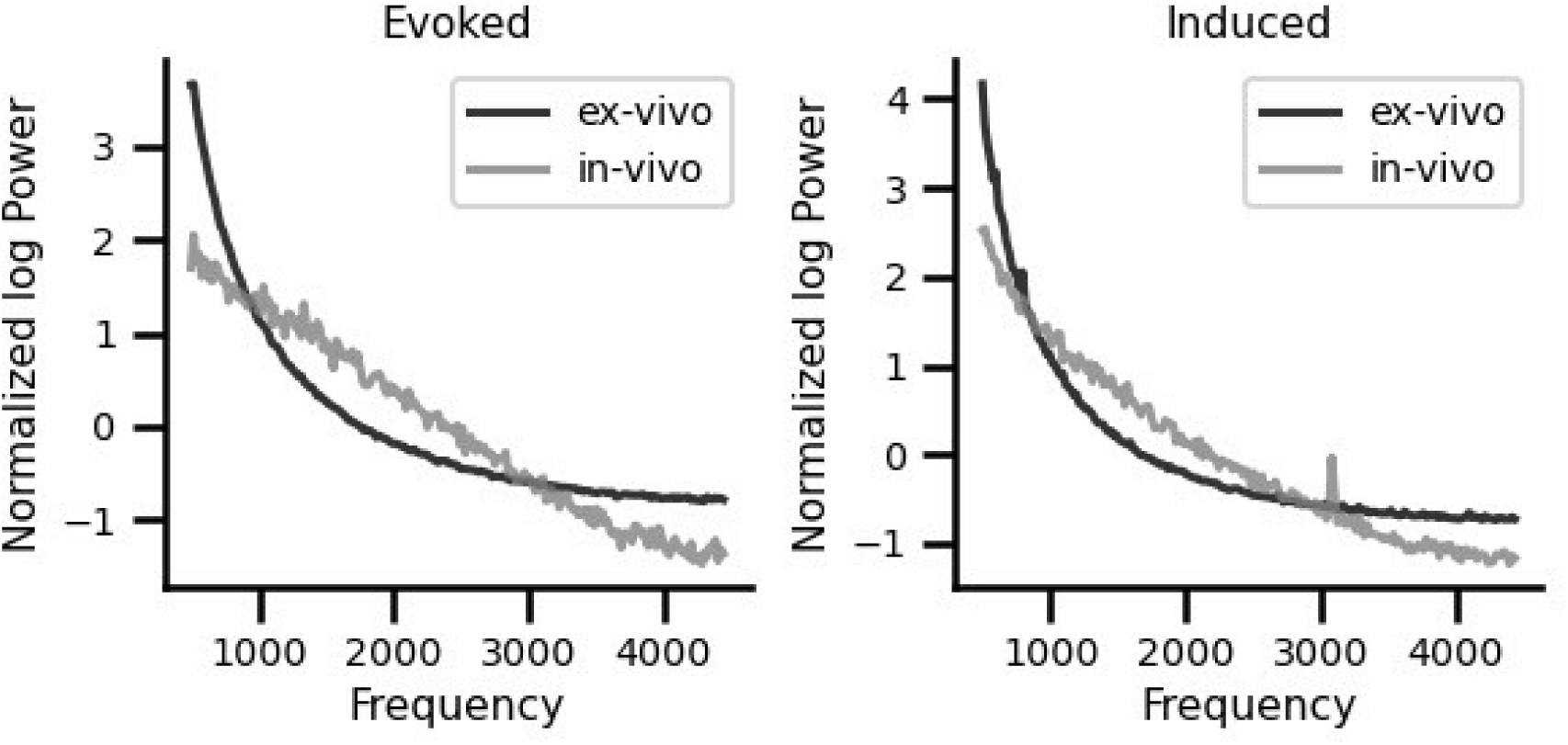
Ex-vivo (black trace) and in-vivo (gray trace) normalized STFT calculated from data analyzed with evoked and induced frequency-analysis methods in spontaneous (silence) conditions. There are no peaks in the ex-vivo and in-vivo evoked conditions. Note that, in the cases of induced analyses, there is a peak at 3.1 kHz in the in-vivo condition (gray trace), which is absent in the case of an ex-vivo recording performed with a cochlear implant immersed in saline solution (black trace).

To rule out a possible artifact from the neighboring equipment in the recording rooms, we also performed the experimental protocol in two subjects, in two different soundproof rooms, located in the first and fourth floors of our hospital, obtaining the 3.1 kHz in both rooms (Figure 6). In addition, to rule out a possible auditory stimulation at 3.1 kHz, we performed an additional control using a microphone during the 2 kHz protocol and found no acoustic peak at the 3.1 kHz frequency region (Figure 7).

**Figure 6.**
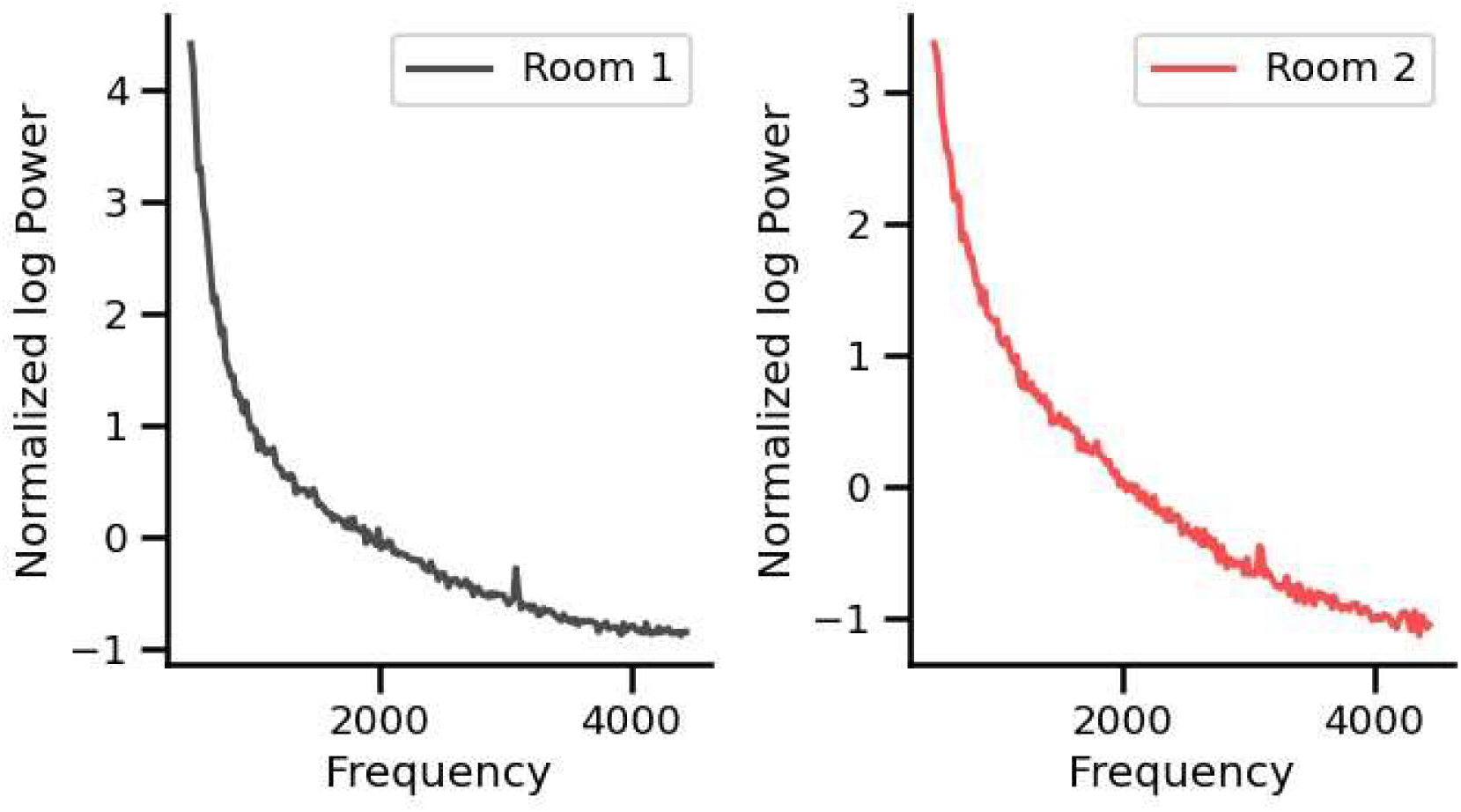
Normalized power spectrums of the cochlear implant noise obtained from one volunteer in two different rooms located in the first and fourth floors in the hospital during silent conditions. Induced FFTs were normalized to the aperiodic 1/f baseline curve. Note that the 3.1 kHz frequency peak is present in both rooms.

**Figure 7.**
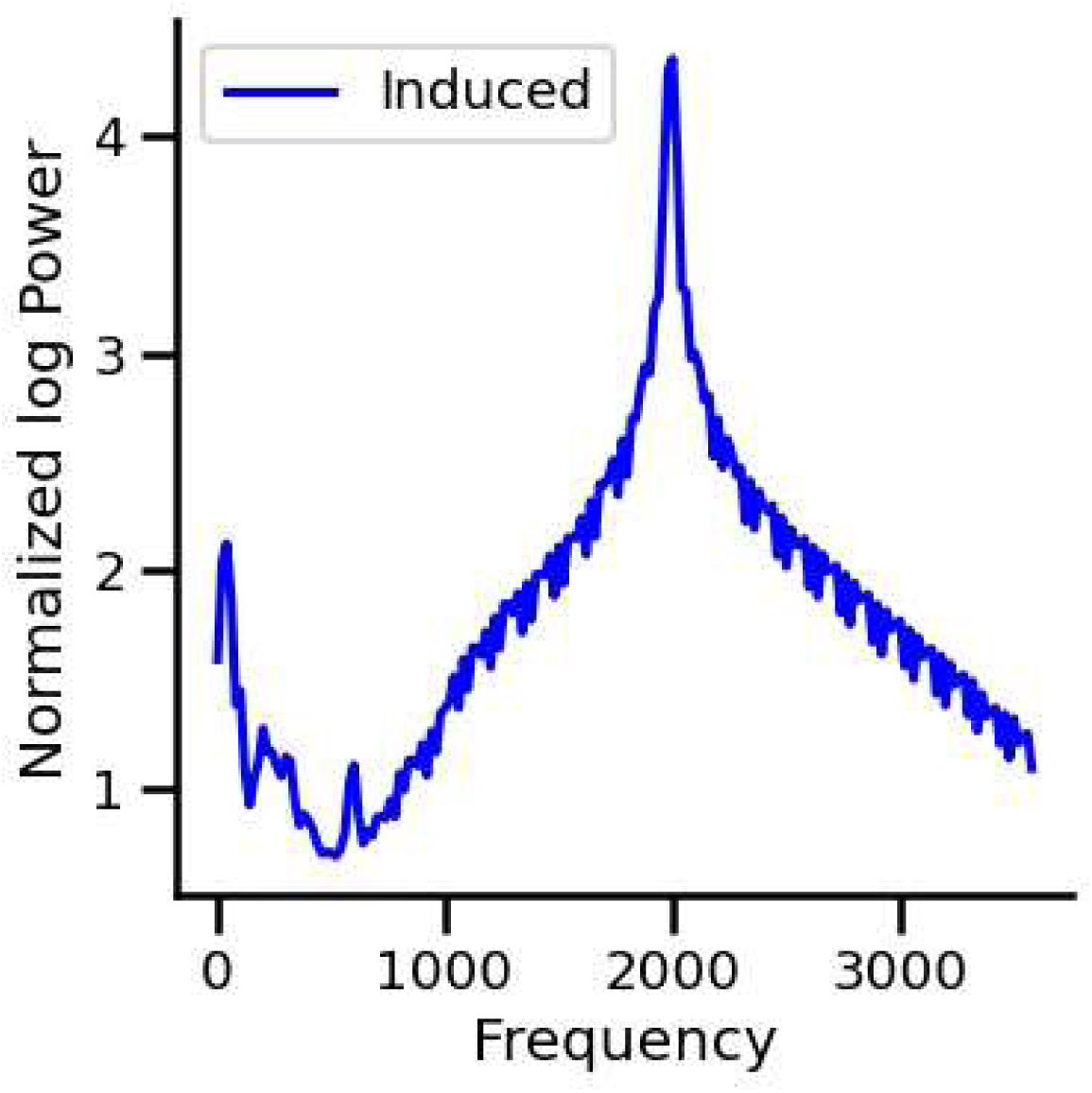
Microphone control using the induced analysis method to compute STFT to the acoustic signal. There is no acoustic peak at 3.1 kHz during the 2 kHz stimulation of the experimental protocol.

Next, we evaluated the number of trials needed to obtain the two types of spectrum noises (peak at 3.1 kHz and the oscillatory pattern) and found that ∼100 trials are necessary to obtain a reliable peak at 3.1 kHz (Figure 8), while the oscillatory pattern spectrum appears with less than 10 trials. To control for possible aliasing artifacts, we randomly resampled the signal into a new sampling frequency between 7 and 10 kHz, and repeated this 1000 times. We found no difference in the 3.1 kHz peak. Additionally, we oversampled the signal up to 10 times Nyquist frequency, to relax anti-aliasing filters and possible phase distortion of the signal, and found no effect in the detection of 3.1 kHz peak (Figure 9).

**Figure 8.**
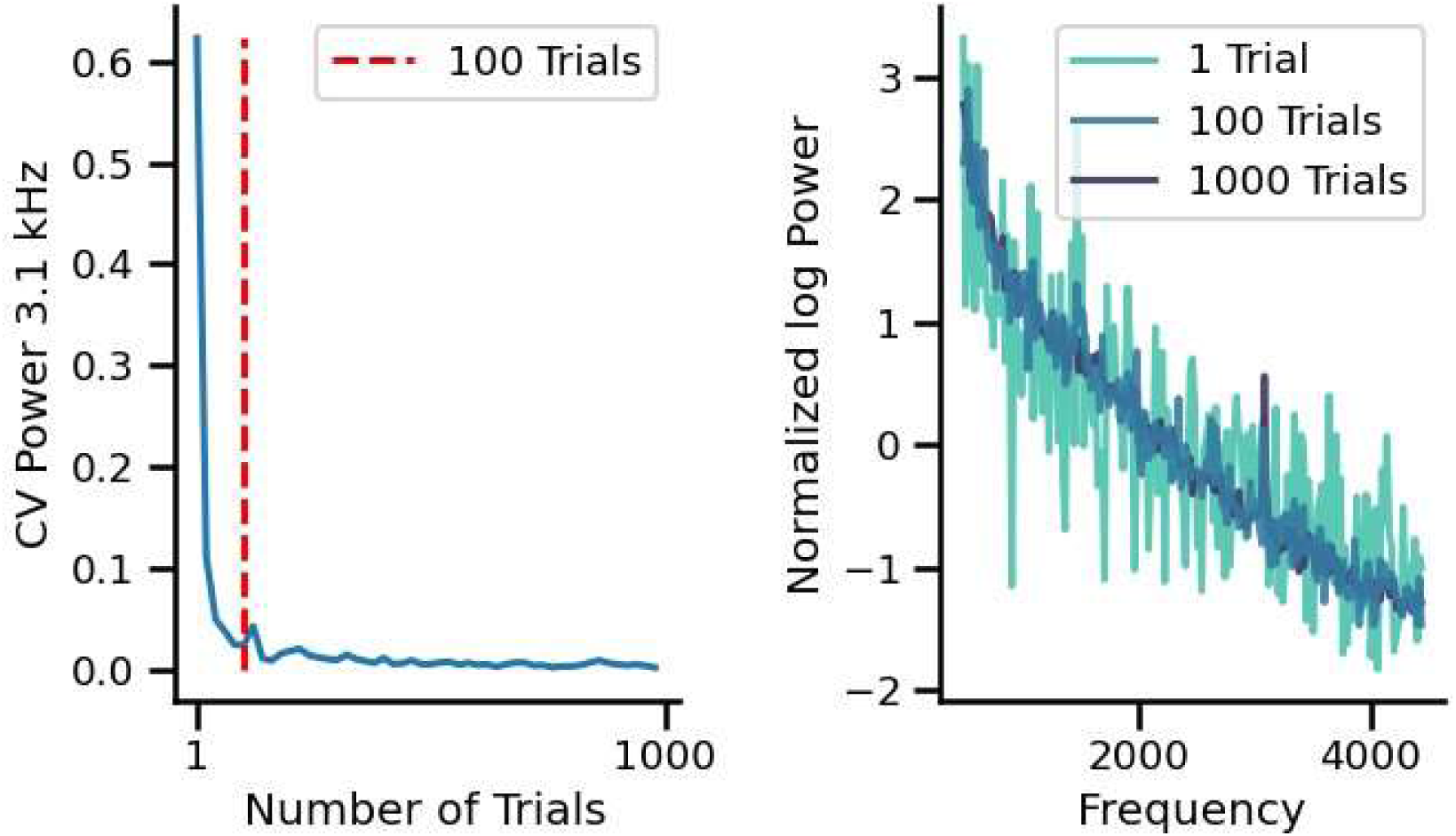
Number of trials needed for obtaining a reliable 3.1 kHz peak. Left panel shows that at 100 trials the coefficient of variation of the amplitude of the 3.1 kHz peak reaches an asymptotic value (red segmented vertical line). Right panel illustrates the power spectrum of the data analyzing 1, 100 and 1000 trials, showing that a minimum of 100 trials is necessary for obtaining a reliable peak at 3.1 kHz.

**Figure 9.**
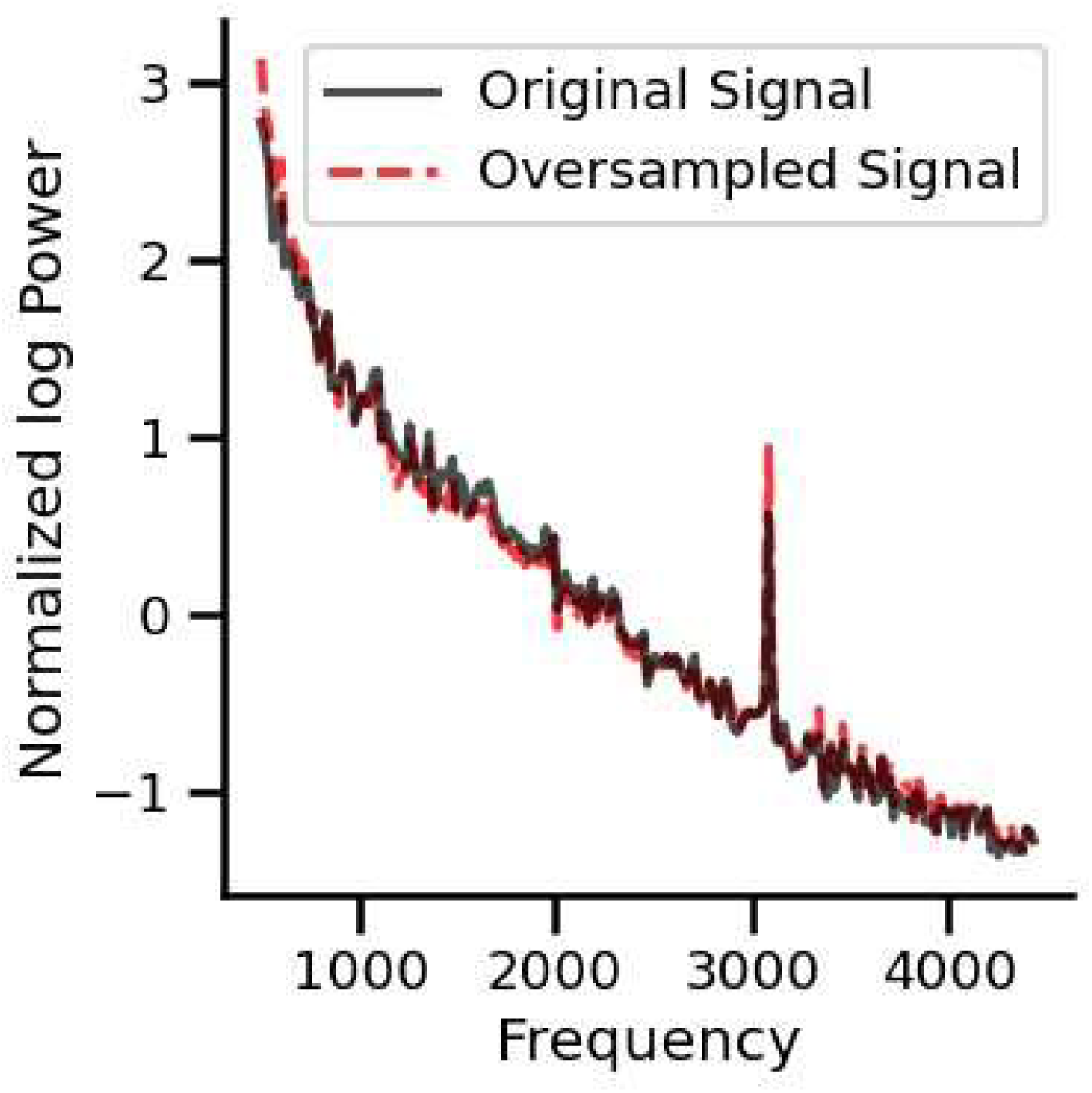
Normalized power spectra obtained in silent conditions using oversampling methods. After reconstructing the signal, the frequency peak at 3.1 kHz remained stable.

Finally, we evaluated whether the amplitude of the 3.1 kHz frequency peak was affected by the auditory 0.5 and 2 kHz stimuli in the 11 cases that displayed the 3.1 kHz frequency peak in the induced activity. There were non-significant changes in the amplitude of the 3.1 kHz peak with 0.5 kHz and 2 kHz tones (Figure 10).

**Figure 10.**
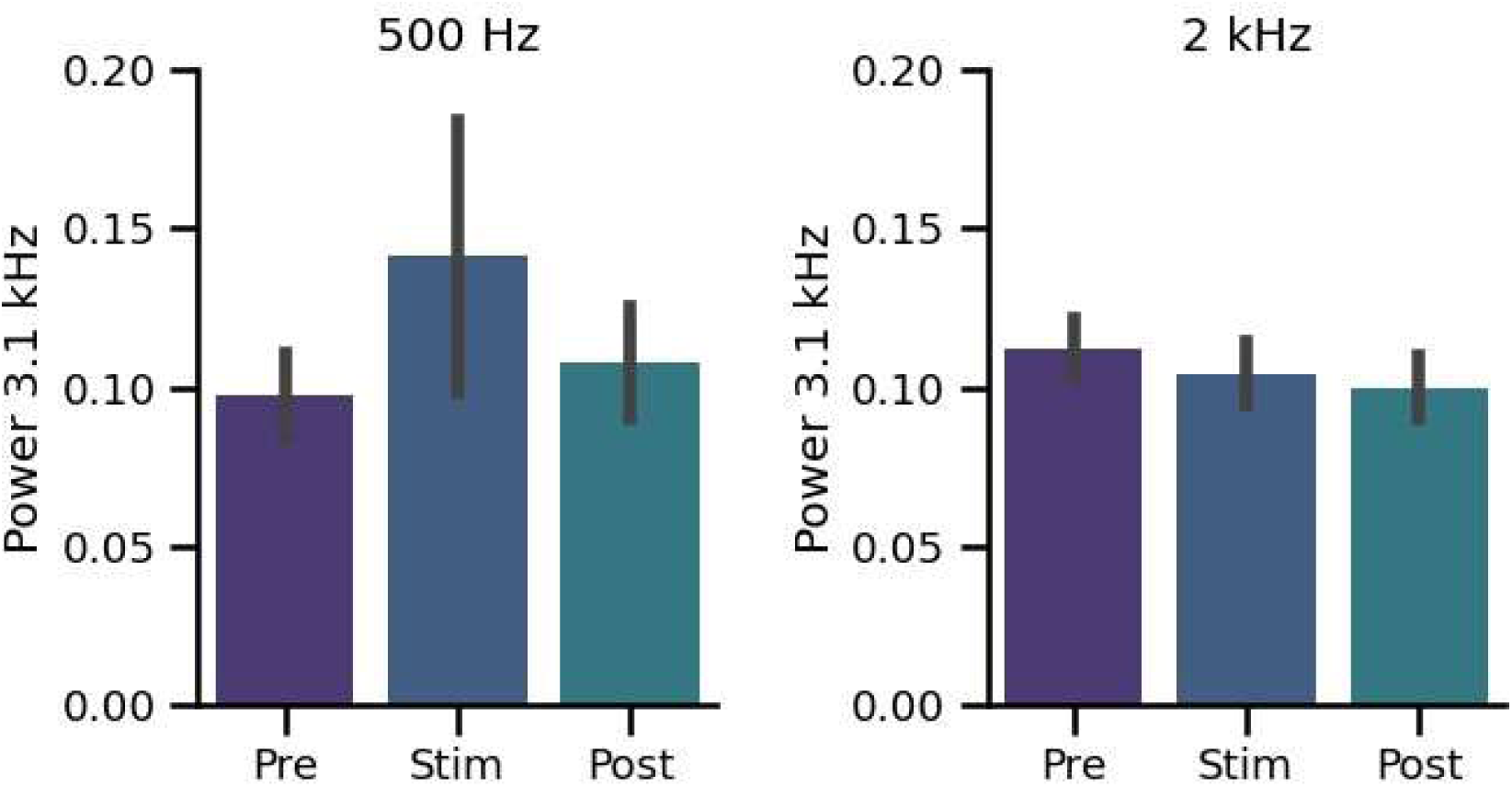
Grand average of the amplitude of the frequency peak at 3.1 kHz in the three periods of the nine minutes protocol (purple: baseline (Pre), blue: during 0.5 kHz and 2 kHz stimulation, cyan: post-stimulation). We found a non-significant increase of the amplitudes of the 3.1 kHz peak with 0.5 kHz and 2 kHz tones.

## Discussion

We found a frequency peak at 3.1 kHz in the electrocochleography spectrum obtained by averaging single-trials STFT during spontaneous (silent) and stimulus-induced conditions in 55% of the cochlear implant subjects. In addition, in 35% of the subjects, we found an oscillatory pattern that masked the power spectrum at the 3.1 kHz frequency band.

## A biological origin for the 3.1 kHz peak in the electrocochleography noise of cochlear implants

The discovery of a new frequency component at 3.1 kHz recorded from the single-trial noise of the cochlear implant electrocochleography raises several questions, including whether this component arises from a biological source. Several controls were performed to try to answer this question: (1) the peak is not present when recording from an ex-vivo cochlear implant in saline solution; (2) in our experimental setup there is no auditory stimulus at 3.1 kHz as evidenced by the microphone control; (3) re-sampling methods are able to reconstruct the signal and re-obtain the peak at 3.1 kHz; (4) there is a need of averaging at least ∼100 trials to improve the signal to noise ratio; (5) the 3.1 kHz peak was obtained in different soundproof rooms of the hospital, one located in the fourth floor and the other in the first floor; (6) The fact that the 3.1 kHz frequency peak was observed only when using induced analysis methods, but not with evoked analysis, indicates that the origin of this signal has temporal jitter, which is generally observed in biological signals. Taken together, these arguments let us propose that the 3.1 kHz frequency peak obtained from the cochlear implant electrocochleography in spontaneous conditions, probably has a biological origin.

On the other hand, the oscillatory pattern observed in around one third of the subjects (7/20) is probably an artifact, as it has multiple frequency components that are evident with less than 10 averaged trials. Understanding the origin of this oscillatory noise could help to develop tools to remove this oscillatory noise and clean the spectrum signal, which might probably help in obtaining the 3.1 kHz frequency peak.

## Is the 3.1 kHz frequency peak a measure of spontaneous auditory-nerve activity?

We have argued that the 3.1 kHz frequency peak probably has a biological origin. The next question is whether it is a measure of the spontaneous auditory-nerve activity. The amplitude of the 3.1 kHz frequency peak did not change with pure tone stimulation at 0.5 and 2 kHz. These results are intriguing, but they could be explained by the fact that we were using pure tones to induce auditory responses in severe to profound deaf subjects. In addition, the 3.1 kHz frequency peak needs hundreds of single-trials to emerge from background noise. We can speculate, in a putative scenario, that with better residual hearing and using wide band stimuli, such as white noise or clicks as auditory elicitors, it might be possible to obtain significant changes in the amplitude of the 3.1 kHz frequency peak.

One interesting argument in favor of single unit origin for the 3.1 kHz peak is the temporal profile of auditory-nerve neuron waveforms. The waveform of auditory-nerve neurons in humans is unknown, while different animal models have given insights into their temporal profile (Kiang et al., 1976; Manley et al., 1976; Liberman, 1978; Wang, 1979; Temchin et al., 2008). These works show that single unitary activity of the auditory-nerve neurons in animals lasts less than 1 ms.

Regarding humans, deconvolution algorithms have been used to predict single fiber unitary waveforms from ECAPs recorded with electrocochleography in cochlear implant users (Dong et al., 2020; Dong et al., 2021; Dong et al., 2023). These works have predicted a width for auditory-nerve single fiber waveforms of around 0.2 to 0.4 ms, which is in agreement with a peak at 3.1 kHz (predicting a waveform width of 0.32 ms in humans). However, whether this frequency peak at 3.1 kHz corresponds to a measure of single fiber activity of the auditory-nerve is still an open question.

## Induced versus evoked analyses in electrocochleography

The standard methods for obtaining cochlear microphonics with frequency analyses in electrocochleography calculate the fast-Fourier transform of the averaged waveform in time, corresponding to an “evoked” method (Koka et al., 2017; Giardina et al., 2019;

ÓLeary et al., 2023). From animal studies, it is known that the evoked potential analysis methods can eliminate important information from the single-trial frequency domain that is not time-locked to the synchronizing stimuli (Delano et al., 2008; Yusuf et al., 2017). In our experiments, we found that the cochlear microphonic responses (to the 0.5 and 2 kHz stimuli) of the same data were larger when evaluated with evoked methods than with induced methods. On the other hand, the 3.1 kHz frequency peak only appeared when using induced methods. Therefore, we propose the combined use of evoked and induced methods for obtaining complimentary information from the cochlear implant electrocochleography signal.

## Conclusion

We found two main types of patterns in the frequency analysis of the single-trial cochlear implant electrocochleography noise, including a frequency peak at 3.1 kHz that might correspond to auditory-nerve spontaneous activity, and an oscillatory pattern that probably reflects an artifact.

## Acknowledgement

Funded by FONDECYT 1220607, FONDECYT 3230557 and Fondo BASAL ANID FB0008. We thank Smita Agrawal and Ana-Claudia Martinho de Carvalho for helpful comments and suggestions.

## Conflict of interest

PHD has a research contract with Advanced Bionics who had no role in the design and experimental procedures of the present work.

## Notes

### Competing Interest Statement

Paul H. Delano has a research contract with Advanced Bionics who had no role in the design and experimental procedures of the present work.

## References

1. Carlyon RP, Goehring T. Cochlear Implant Research and Development in the Twenty-first Century: A Critical Update. J Assoc Res Otolaryngol. 2021;22(5):481–508. doi:10.1007/s10162-021-00811-5

2. Delano PH, Elgueda D, Hamame CM, Robles L. Selective attention to visual stimuli reduces cochlear sensitivity in chinchillas. J Neurosci. 2007;27(15):4146–4153. doi:10.1523/JNEUROSCI.3702-06.2007

3. Delano PH, Pavez E, Robles L, Maldonado PE. Stimulus-dependent oscillations and evoked potentials in chinchilla auditory cortex. J Comp Physiol A Neuroethol Sens Neural Behav Physiol. 2008;194(8):693–700. doi:10.1007/s00359-008-0340-4

4. Dolan DF, Teas DC, Walton JP. Relation between discharges in auditory nerve fibers and the whole-nerve response shown by forward masking: an empirical model for the AP. J Acoust Soc Am. 1983;73(2):580–591. doi:10.1121/1.389005

5. Dolan DF, Nuttall AL, Avinash G. Asynchronous neural activity recorded from the round window. J Acoust Soc Am. 1990;87(6):2621–2627. doi:10.1121/1.399054

6. Dong Y, Briaire JJ, Biesheuvel JD, Stronks HC, Frijns JHM. Unravelling the temporal properties of human eCAPs through an iterative deconvolution model. Hear Res. 2020;395:108037. doi:10.1016/j.heares.2020.108037

7. Dong Y, Stronks HC, Briaire JJ, Frijns JHM. An iterative deconvolution model to extract the temporal firing properties of the auditory nerve fibers in human eCAPs. MethodsX. 2021;8:101240. Published 2021 Jan 22. doi:10.1016/j.mex.2021.101240

8. Dong Y, Briaire JJ, Stronks HC, Frijns JHM. Speech Perception Performance in Cochlear Implant Recipients Correlates to the Number and Synchrony of Excited Auditory Nerve Fibers Derived From Electrically Evoked Compound Action Potentials. Ear Hear. 2023;44(2):276–286. doi:10.1097/AUD.0000000000001279

9. Donoghue T, Haller M, Peterson EJ, et al. Parameterizing neural power spectra into periodic and aperiodic components. Nat Neurosci. 2020;23(12):1655–1665. doi:10.1038/s41593-020-00744-x

10. Eggermont JJ. Cochlea and auditory nerve. Handb Clin Neurol. 2019;160:437-449. doi:10.1016/B978-0-444-64032-1.00029-1

11. Galambos R. Suppression of auditory nerve activity by stimulation of efferent fibers to cochlea. J Neurophysiol. 1956;19(5):424–437. doi:10.1152/jn.1956.19.5.424

12. Giardina CK, Brown KD, Adunka OF, et al. Intracochlear Electrocochleography: Response Patterns During Cochlear Implantation and Hearing Preservation. Ear Hear. 2019;40(4):833–848. doi:10.1097/AUD.0000000000000659

13. Hey M, Müller-Deile J. Accuracy of measurement in electrically evoked compound action potentials. J Neurosci Methods. 2015;239:214–222. doi:10.1016/j.jneumeth.2014.10.012

14. Ishikawa M, Kojima A, Terao S, Nagai M, Kusaka G, Naritaka H. Cochlear Nerve Action Potential Monitoring for Preserving Function of an Unseen Cochlear Nerve in Vestibular Schwannoma Surgery. World Neurosurg. 2017;106:1057.e1–1057.e7. doi:10.1016/j.wneu.2017.07.113

15. Kiang NY. Discharge patterns of single fibers in the cat’s auditory nerve. MASSACHUSETTS INST OF TECH CAMBRIDGE RESEARCH LAB OF ELECTRONICS; 1965.

16. Kiang NY, Sachs MB, Peake WT. Shapes of tuning curves for single auditory-nerve fibers. J Acoust Soc Am. 1967;42(6):1341–1342. doi:10.1121/1.1910723

17. Kiang NY, Liberman MC, Levine RA. Auditory-nerve activity in cats exposed to ototoxic drugs and high-intensity sounds. Ann Otol Rhinol Laryngol. 1976;85(6 PT. 1):752–768. doi:10.1177/000348947608500605

18. Koka K, Saoji AA, Litvak LM. Electrocochleography in Cochlear Implant Recipients With Residual Hearing: Comparison With Audiometric Thresholds. Ear Hear. 2017;38(3):e161–e167. doi:10.1097/AUD.0000000000000385

19. Kujawa SG, Liberman MC. Adding insult to injury: cochlear nerve degeneration after “temporary” noise-induced hearing loss. J Neurosci. 2009;29(45):14077–14085. doi:10.1523/JNEUROSCI.2845-09.2009

20. Liberman MC. Auditory-nerve response from cats raised in a low-noise chamber. J Acoust Soc Am. 1978;63(2):442–455. doi:10.1121/1.381736

21. Louage DH, van der Heijden M, Joris PX. Temporal properties of responses to broadband noise in the auditory nerve. J Neurophysiol. 2004;91(5):2051–2065. doi:10.1152/jn.00816.2003

22. Maggu AR. Auditory Evoked Potentials in Communication Disorders: An Overview of Past, Present, and Future. Semin Hear. 2022;43(3):137–148. Published 2022 Oct 26. doi:10.1055/s-0042-1756160

23. Manley GA, Robertson D. Analysis of spontaneous activity of auditory neurones in the spiral ganglion of the guinea-pig cochlea. J Physiol. 1976;258(2):323–336. doi:10.1113/jphysiol.1976.sp011422

24. O’Leary S, Mylanus E, Venail F, et al. Monitoring Cochlear Health With Intracochlear Electrocochleography During Cochlear Implantation: Findings From an International Clinical Investigation. Ear Hear. 2023;44(2):358–370. doi:10.1097/AUD.0000000000001288

25. Pardo-Jadue J, Dragicevic CD, Bowen M, Delano PH. On the Origin of the 1,000 Hz Peak in the Spectrum of the Human Tympanic Electrical Noise. Front Neurosci. 2017;11:395. Published 2017 Jul 11. doi:10.3389/fnins.2017.00395

26. Rauschecker JP, Shannon RV. Sending sound to the brain. Science. 2002;295(5557):1025–1029. doi:10.1126/science.1067796

27. Searchfield GD, Muñoz DJ, Thorne PR. Ensemble spontaneous activity in the guinea-pig cochlear nerve. Hear Res. 2004;192(1-2):23–35. doi:10.1016/j.heares.2004.02.006

28. Steinhoff HJ, Böhnke F, Janssen T. Click ABR intensity-latency characteristics in diagnosing conductive and cochlear hearing losses. Arch Otorhinolaryngol. 1988;245(5):259–265. doi:10.1007/BF00464627

29. Temchin AN, Rich NC, Ruggero MA. Threshold tuning curves of chinchilla auditory nerve fibers. II. Dependence on spontaneous activity and relation to cochlear nonlinearity. J Neurophysiol. 2008;100(5):2899–2906. doi:10.1152/jn.90639.2008

30. Undurraga JA, van Wieringen A, Carlyon RP, Macherey O, Wouters J. Polarity effects on neural responses of the electrically stimulated auditory nerve at different cochlear sites. Hear Res. 2010;269(1-2):146–161. doi:10.1016/j.heares.2010.06.017

31. Yamakami I, Oka N, Yamaura A. Intraoperative monitoring of cochlear nerve compound action potential in cerebellopontine angle tumour removal. J Clin Neurosci. 2003;10(5):567–570. doi:10.1016/s0967-5868(03)00143-7

32. Yusuf PA, Hubka P, Tillein J, Kral A. Induced cortical responses require developmental sensory experience. Brain. 2017;140(12):3153–3165. doi:10.1093/brain/awx286

33. Wang B. The relation between the compound action potential and unit discharges of the auditory nerve (Doctoral dissertation, Massachusetts Institute of Technology), 1979.

